# Release probability increases towards distal dendrites boosting high-frequency signal transfer

**DOI:** 10.1101/2020.08.29.273227

**Authors:** Thomas P. Jensen, Olga Kopach, Leonid P. Savchenko, James P. Reynolds, Dmitri A. Rusakov

**Author notes:** Correspondence: Dmitri Rusakov or Thomas Jensen.

## Abstract

Dendritic integration of synaptic inputs entangles their increased electrotonic attenuation at distal dendrites, which can be counterbalanced by the increased synaptic receptor density. However, during sustained network activity the influence of individual synapses depends on their release properties. How these properties are distributed along dendrites remains poorly understood. Here, we employed classical optical quantal analyses and a genetically encoded optical glutamate sensor in acute hippocampal slices to monitor release at CA3-CA1 synapses. We find that their release probability increases with greater distances from the soma. Similar-fidelity synapses tend to group together whereas release probability shows no trends regarding the within-branch position. Simulations with a realistic CA1 pyramidal cell hosting stochastic synapses suggest that the observed trends boost signal transfer fidelity, particularly at higher input frequencies. Because high-frequency bursting has been associated with learning, the release probability pattern we have found may play a key role in memory trace formation.

## INTRODUCTION

Information processing in brain circuits relies on dendritic integration of excitatory inputs that occur at varied distances from the soma. Whilst the cable impedance of dendrites attenuates electric signals arriving from distal synapses, some nerve cells appear well equipped to counter this trend. In hippocampal CA1 pyramidal neurons, synaptic receptor numbers increase monotonically with the synapse-soma distance (Andrasfalvy and Magee, 2001; Nicholson et al., 2006). This, combined with the regenerative properties of local ion channels (Cash and Yuste, 1999; Magee, 1999), can provide efficient, relatively homogeneous summation of synaptic inputs across the dendritic tree of these cells (Cash and Yuste, 1999; Magee, 2000; Magee and Cook, 2000). In contrast, distal synapses on cortical L5 pyramidal neurons appear relatively ineffective in somatic excitation (Williams and Stuart, 2002). A combination of 3D electron microscopy with biophysical modelling of CA1 pyramidal cells has also argued that the greater occurrence of excitatory synapses towards the dendritic branch origin helps to normalise input efficacy along the dendrites (Katz et al., 2009). However, it remains largely unknown whether the other fundamental synaptic property, neurotransmitter release probability (P_r_) remains constant along the dendrites. Because P_r_ defines how much information synaptic connections transmit over time (Zador, 1998), its distribution pattern is critical to the rules of dendritic input integration.

A recent elegant study has combined electron microscopy with extracellular (paired-pulse) afferent activation of multiple synapses in basal dendrites of CA1 pyramidal cells, to find the average P_r_ values decreasing towards the cell periphery (Grillo et al., 2018). Thus, a few proximal connections appear to have a greater impact on the cell output than more numerous synapses occurring distally. This observation raised the question of the synaptic connectome efficiency, suggesting that, during continued afferent spiking, distal inputs are weakened. Although highly supra-linear local events, such as dendritic spikes could, in principle, compensate for this trend (Larkum and Nevian, 2008; Branco and Hausser, 2009), how prevalent such events are remains debatable. Furthermore, if P_r_ tends to decrease towards more distal apical dendrites in CA1 pyramidal cells, the suggested input summation rules in these cells (Magee, 2000; Magee and Cook, 2000; Branco and Hausser, 2011), and at least in some other principal neurons (Araya et al., 2006), would be broken during sustained network activity.

Methodologically, paired-pulse recordings of multi-synaptic responses (Grillo et al., 2018), while revealing the trend, do not provide direct readout of either P_r_, paired-pulse ratios (PPRs), or the P_r_ distribution among individual connections. This makes it difficult to evaluate how these properties of P_r_ affect dendritic integration. To address these challenges, here we monitored P_r_ in individual CA3-CA1 synapses, by systematically employing single-synapse optical quantal analyses (OQA) (Oertner et al., 2002; Emptage et al., 2003), based on Ca^2+^ imaging in individual dendritic spines, as established previously (Sylantyev et al., 2013; Boddum et al., 2016). In a complementary approach, we employed the genetically encoded optical glutamate sensor iGluSnFR, as established earlier (Jensen et al., 2017; Jensen et al., 2019; Kopach et al., 2020), to document a spatial trend of P_r_ values across the *s. radiatrum*. Exploring the results with a realistic biophysical model of the CA1 pyramidal cell (Migliore et al., 1999) equipped with multiple stochastic synaptic inputs, revealed how the observed trends in the P_r_ pattern along dendrites could affect synaptic input integration in CA1 pyramidal cells.

## RESULTS

### Monitoring release probability with optical quantal analysis

We used 300-350 µm transverse hippocampal slices of the 3-4-week-old rats (Star Methods). First, we held CA1 pyramidal cells in whole-cell (V_m_ = -65 mV) and dialysed them with the red morphological tracer Alexa Fluor 594 (50 µM) and Ca^2+^ sensitive indicators as detailed in Methods and described previously (Sylantyev et al., 2013; Zheng et al., 2015). We visualised cell morphology with two-photon excitation (λ_x_^2p^= 800 nm), and placed the extracellular stimulating pipette in the 10-20 µm proximity of apical dendrites (Figure 1A, Figure S1A). The system was focused on individual dendritic spines that responded to paired-pulse stimuli (50 ms apart) with localised Ca^2+^ transients in a stochastic manner (Figure 1B, Figure S1B). We were thus able to readily distinguish successes and failures of neurotransmitter release. Because individual dendritic spines of CA1 pyramidal cells host almost exclusively only one CA3-CA1 synapse (Harris and Stevens, 1989; Bloss et al., 2018), the Ca^2+^ signal failure counts over 15-30 trials provided direct readout of P_r_ at individual synapses (Figure 1B, Figure S1C), as we showed earlier (Sylantyev et al., 2013; Boddum et al., 2016). Once finished with testing, we recorded the curvilinear distance from the spine of interest to the cell soma (Star Methods).

**Figure 1.**
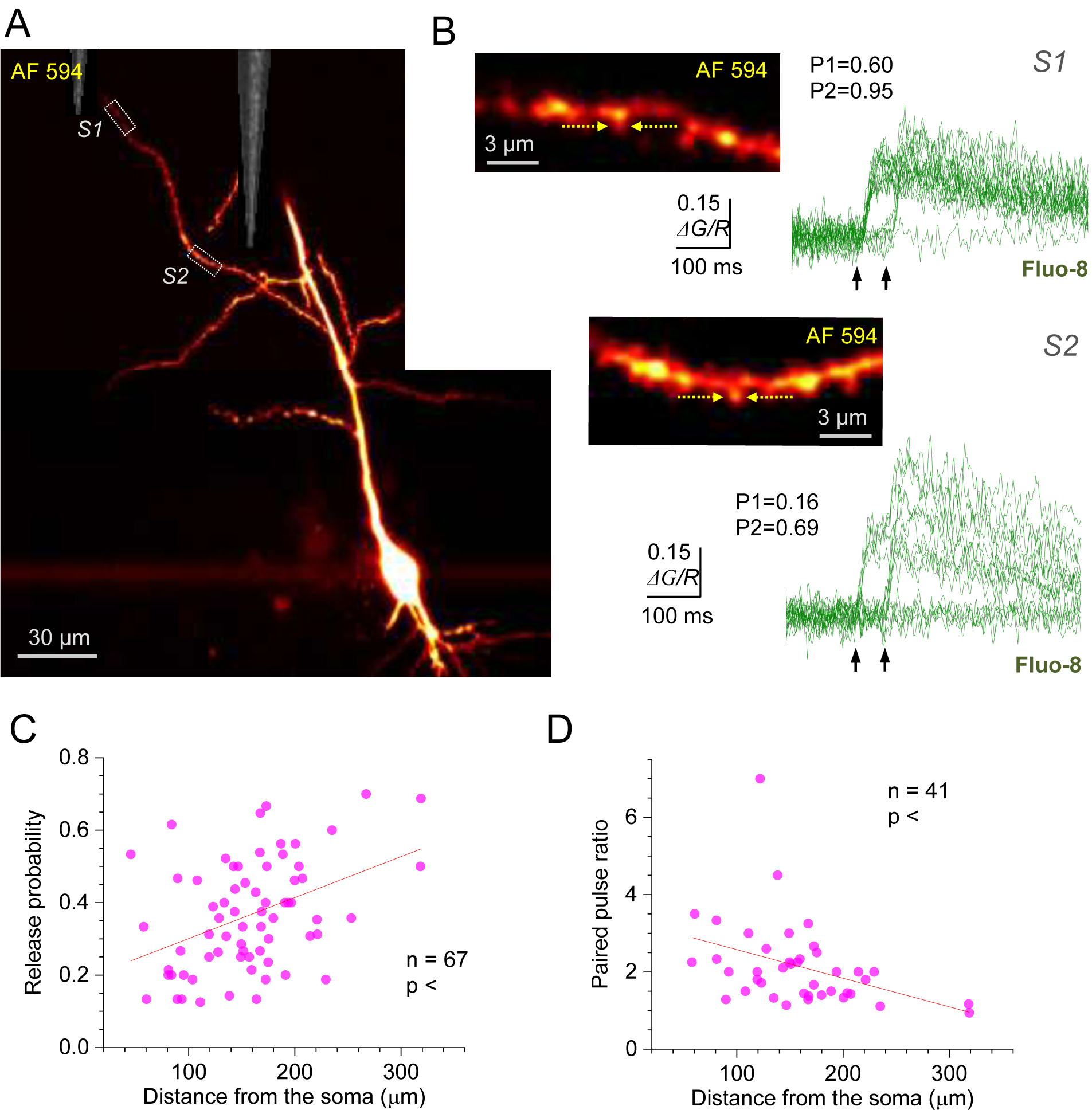
Optical quantal analysis at individual CA3-CA1 synapses reports higher release probability towards distant dendrites. (A) CA1 pyramidal cell held in whole-cell mode (acute hippocampal slice), dialysed with 50 µM AF 594 and 300 µM Fluo-8 (75 µm z-stack average, *λ*_x_^2p^ = 800 nm; AF 594 channel). Dotted rectangles (*S1* and *S2*), two ROIs to record from two dendritic spines; stimulating electrode positions are illustrated by DIC insets. (B) Image panels, ROIs as in (A), shown at higher magnification; arrows, linescan positioning at the dendritic spines of interest. Traces, Ca^2+^ signal (*ΔG/R*, Fluo-8 green-channel signal *ΔG* related to red-channel AF 594 signal *R*) recorded as the width-integrated linescan intensity, at two spines as indicated, in response to paired-pulse afferent stimuli (arrows). Release failures and responses can be clearly separated; P1 and P2, average release probability in response to the first and second stimulus, respectively. (C) Average release probability (P_r_; shown as P1 in B) at individual synapses plotted against distance to the soma. Solid line, linear regression (non-zero slope at p < 0.005, Pearson’s r = 0.413, sample size n as indicated). (D) Paired-pulse ratio (as P2/P1 in B) plotted against distance to the soma. Other notations as in (C) (Pearson’s r = -0.392).

A systematic application of this protocol at n = 67 individual synapses (53 cells) has revealed a highly significant trend towards higher P_r_ at more distal synapses (Figure 1C), with average P_r_ = 0.36 ± 0.02 (mean ± SEM). The linear regression for the data suggested a rise in average P_r_ from ∼0.2 at 50 µm to ∼0.5 at 300 µm (range 0.05-0.7; Figure 1C). Over this distance range, the paired-pulse ratio of release probabilities decreased from ∼3 to ∼1.1 (regression at p < 0.001, n = 41; average PPR 2.14 ± 0.17; range 1-7, Figure 1D).

The centrifugal trend in P_r_ values remained significant when the distance was measured from the primary apical dendrites (Figure 2A), suggesting that this trend applies to oblique branches rather than simply reflecting a position along the main dendrite. Interestingly, the analysis of P_r_ values recorded in pairs of synapses located on the same dendrite has revealed that the connections with similar P_r_ tend to occur close to one another (Figure 2B). Thus, ‘homogenisation’ of P_r_ among local synapses, which was detected previously in cultured neurons (Branco et al., 2008), is likely to occur in ex vivo brain tissue. At the same time, we found no trends in P_r_ values with respect to the synapse position relative to the origin, or the end, of the host dendritic branch (Figure 2C). The prevalence of larger, more densely packed spines near the dendritic branch origins was earlier reported to reflect normalisation of synaptic signals integrated at the soma (Katz et al., 2009). In this context, no gradient in P_r_ values within a branch (Figure 2C) suggests that the above normalisation should still be valid during sustained afferent activity.

**Figure 2.**
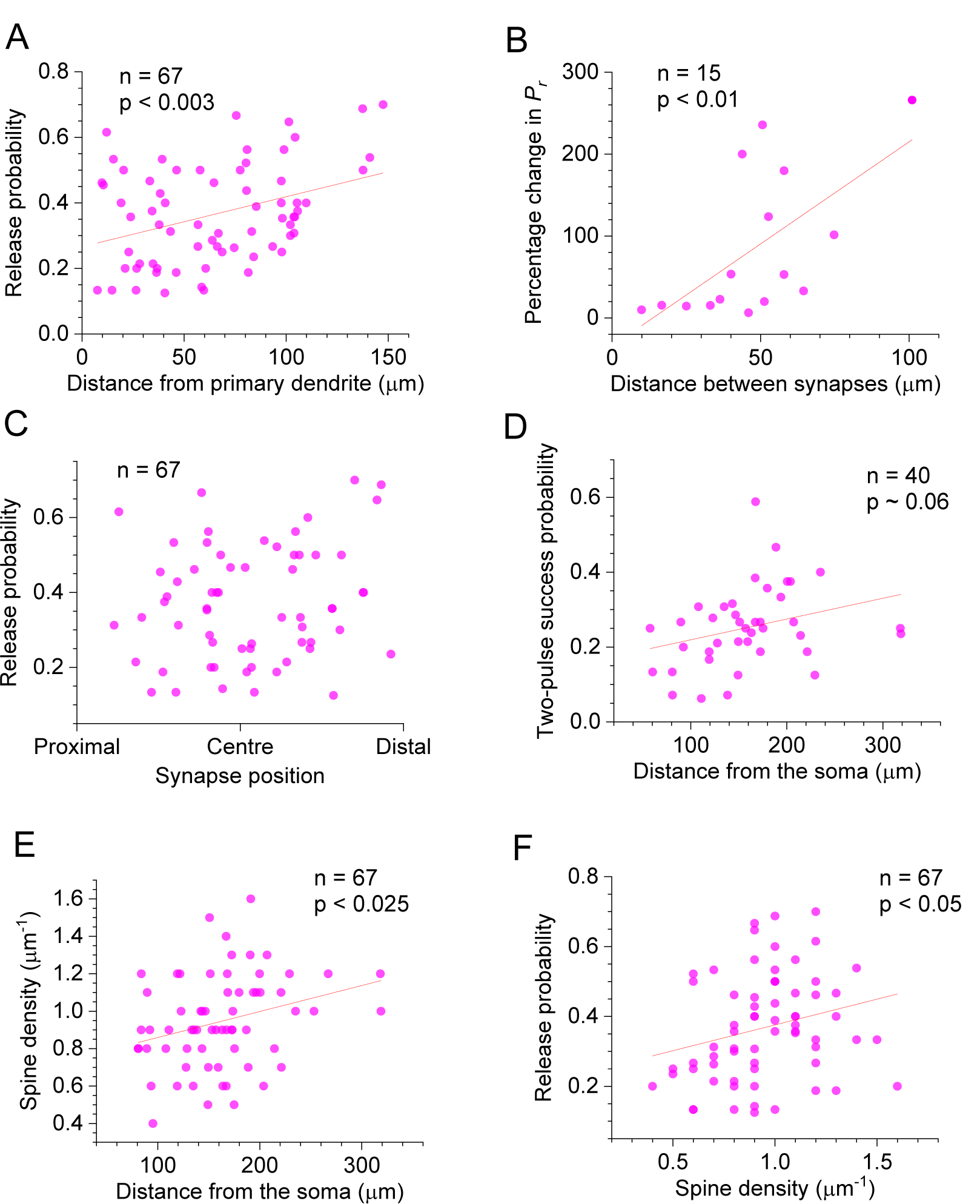
Selected features of excitatory synapses with respect to their dendritic location. (A) Average release probability (P_r_) at individual synapses plotted against distance to the first dendrite branching point. Solid line, linear regression (non-zero slope at p < 0.01; Person’s r = 0.378). (B) Percentage difference in P_r_ between two synapses on one dendritic branch (as in Figure 1A, Figure S1A), plotted against the distance between them along the branch. Other notations as in (A) (Pearson’s r = 0.623). (C) Average release probability (P_r_) plotted against relative synapse position at the dendritic branch: synapse co-ordinate was scaled to the 0-1 range representing the branch origin and the end, as indicated. (D) Probability of release success upon both afferent stimuli, plotted against distance from the soma. Other notations as in (A) (Pearson’s r = 0.300). (E) Apparent spine density along the dendrite (smallest/thinnest spines could be undetectable, see Discussion), plotted against distance from the soma. Other notations as in (A) (Pearson’s r = 0.286). (F) Release probability (P_r_), plotted against distance from the soma. Other notations as in (A) (Pearson’s r = 0.245).

Because higher P_r_ are normally associated with lower PPRs (as in Figure 1D), the probability of both stimuli initiating successful release showed only a barely detectable increase with the spine-soma distance (Figure 2D) whereas the second-release P_r_ values were evenly spread along the dendrites (Figure S2A). However, the longitudinal density of optically identifiable dendritic spines increased towards distal dendrites (Figure 2E), also showing positive correlation with the P_r_ (Figure 2F), but no correlation with the PPR values (Figure S2B).

### Monitoring glutamate release at CA3-CA1 synapses with an optical sensor

We have previously shown (Jensen et al., 2017; Jensen et al., 2019) that the patterns of release probability at CA3-CA1 synapses can be successfully gauged by monitoring presynaptic glutamate release with the genetically encoded optical sensor iGluSnFR (Marvin et al., 2018). We therefore employed this method to validate our conclusions obtained with the classical OQA described above. Once iGluSnFR (AAV9.hSynap.iGluSnFR.WPRE.SV40) was expressed in the area CA1 *s*.*radiatum*, we focused on individual presynaptic boutons that showed a fluorescence response to single or paired-pulse stimuli applied to Schaffer collaterals (Methods); the distance between the bouton and the *s. pyramidale* border was documented (Figure 3A). We thus recorded fluorescence responses to paired-pulse stimuli (50 ms apart) using a high-resolution spiral linescan (‘Tornado’) mode, as described earlier (Jensen et al., 2017; Jensen et al., 2019) (Figure 3B, image panels). In such recordings, the high-affinity iGluSnFR can detect glutamate transients at up to ∼1 µm from its synaptic release site (Jensen et al., 2019). Because synapses in area CA1 are packed at a density of ∼2 µm^3^ (or ∼0.5 µm apart) (Rusakov and Kullmann, 1998), in our experiments the iGluSnFR response should detect glutamate molecules released from the immediate synapse, and the residual glutamate escaping from neighbours that happen to be activated by extracellular stimulation. Consequently, we rarely detected release failures in such recordings (over ∼20 trials; Figure 3B, traces), which prevented direct estimation of P_r_ at individual synapses. However, this approach provided robust estimates of the PPR values for glutamate release at (small groups of) synaptic connections occurring at different distances from the CA1 pyramidal cell body layer (PPR, mean ± SEM: 1.49 ± 0.01, n = 33; Figure 3B). The PPR-distance relationship revealed a clear trend towards lower PPR values, hence higher P_r_ values, with greater distances from the *s. pyramidale* (regression at p < 0.001, n = 33; Figure 3C). This was qualitatively consistent with the PPR data obtained with optical quantal analysis of Ca^2+^ signals (Figure 1D). In contrast, the first iGluSnFR response amplitude on its own did not show any significant distance dependence (*ΔF/F*_0,_ mean ± SEM: 37 ± 0.5%; range 0.18-0.81; Figure 3D). This was expected because, in addition to P_r_ per se, the iGluSnFR signal amplitude depends on several poorly controlled local concomitants, such as the degree of glutamate spillover and/or density of activated axons.

**Figure 3.**
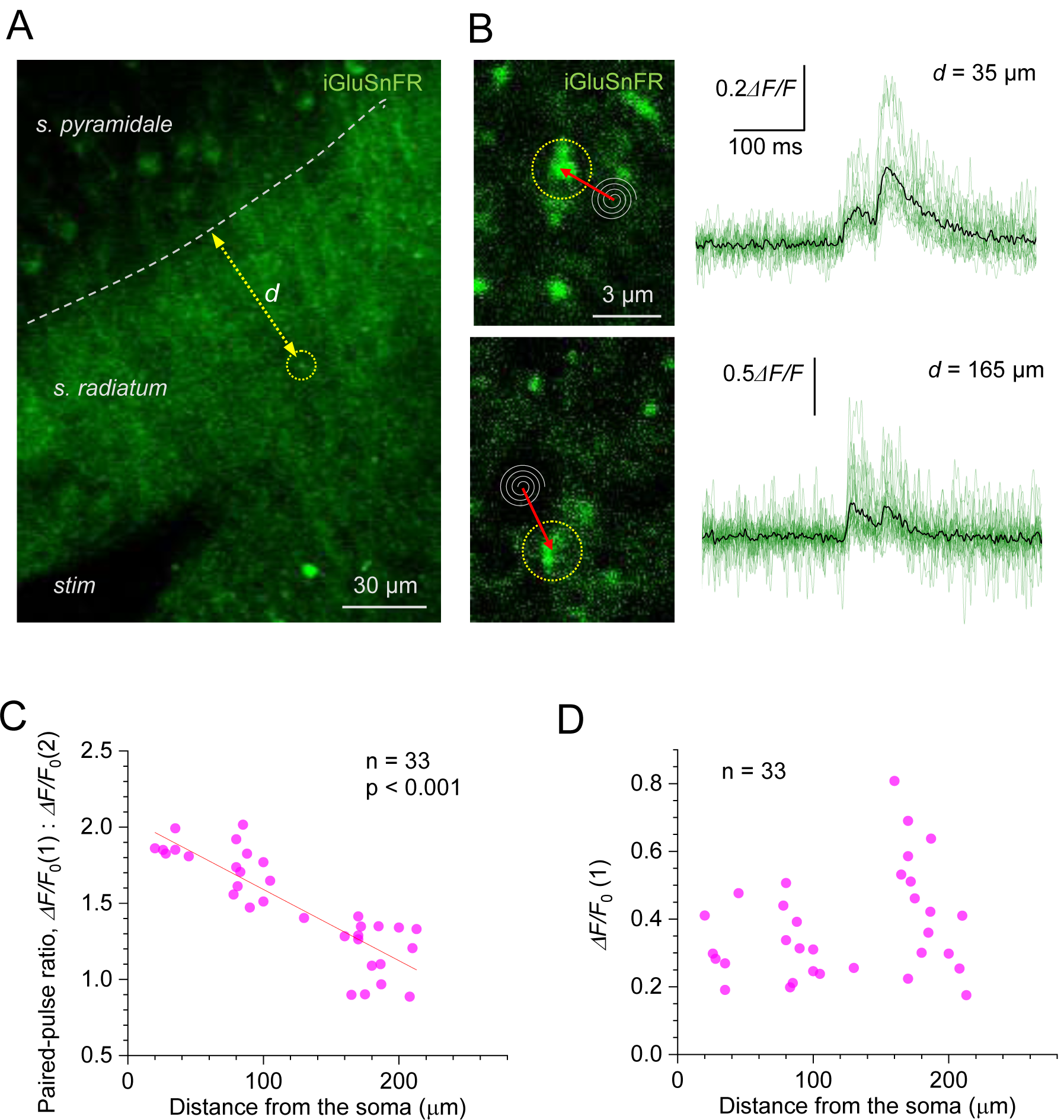
Evoked glutamate release from Schaffer collaterals show lower paired-pulse ratios at greater distances from pyramidal cell bodies. (A) Experimental design: area of the hippocampal slice with iGluSnFR expressed in neuronal membranes (green channel); arrow, measured distance *d* between CA1 pyramidal cell body layer and the axonal bouton of interest; *stim*, stimulating electrode. (B) Examples of recorded axonal boutons (image panels, dotted circles; position of spiral ‘Tornado’ linescans is illustrated) showing characteristic glutamate signals in response to two afferent stimuli 50 ms apart (green traces, individual trials; black, average), at two distances from the *s. pyramidale*, as indicated. The lack of release failures reflects detection of glutamate escaping from multiple neighbouring synapses. (C) Paired-pulse ratio for optical glutamate signals: *ΔF/F*_0_(1) / *ΔF/F*_0_(2) averaged over 18-36 trials at individual boutons, plotted against distance to the soma. Other notations as in Figure 1C (Pearson’s r = -0.863). (D) Amplitude of the first glutamate response, *ΔF/F*_0_(1) averaged over 18-36 trials at individual boutons, plotted against distance to the soma. The amplitude values reflect the average amount of glutamate released from the bouton of interest, and glutamate escaping from its neighbours; other notations as in C.

In a complementary approach, we recorded iGluSnFR responses to a short burst of five afferent stimuli (at 20 Hz), in an attempt to relate their use-dependent release properties to their location with respect to the *s. pyramidale*. To optimise ROI selection under burst stimulation, these recordings were carried out in a frame (time-lapse) mode, as detailed previously (Kopach et al., 2020) (Figure S3A). While such recordings have relatively low temporal resolution, the underlying (fast) fluorescence kinetics could be reconstructed using a straightforward fitting procedure (Methods; Figure S3B). We used the slope of the five-pulse *ΔF/F*_*0*_ signal between the onsets of 1st and 5th stimuli (Figure S3B) as an indicator of short-term release plasticity during the burst. Intriguingly, the slope values increased significantly with greater distances to the *s. pyramidale* (Figure S3C), suggesting greater fidelity in signal transfer by spike bursts towards more distal dendrites, as further explored below.

### A realistic biophysical model explains the role of the release probability trend

To understand how the uneven distribution of P_r_ values affects signal handling by the postsynaptic cell, an earlier study used simulations with a sphere-and-cylinder cell model (Grillo et al., 2018). Here, we employed a realistic multi-compartmental model of a reconstructed CA1 pyramidal cell (Migliore et al., 1999), with 50 excitatory synapses distributed along apical dendrites, so that their positions, density, and P_r_ could be set as required (Figure 4A, Star Methods). In the first test, we asked whether and how the documented centrifugal increase in P_r_ and synaptic density (termed ‘P_r_ trend’ and ‘density trend’, respectively) changes cell output under a synchronous discharge of Schaffer collaterals. A series of 100 paired-pulse stimulus test runs was simulated, each generating stochastic ‘glutamate release’ at the 50 synapses, either with the same average P_r_ (0.36) at uniformly scattered synapses, or with the P_r_ values distributed in accord with our data (Figure 1C), or, additionally, with the synaptic density trend as found (Figure 2E).

**Figure 4.**
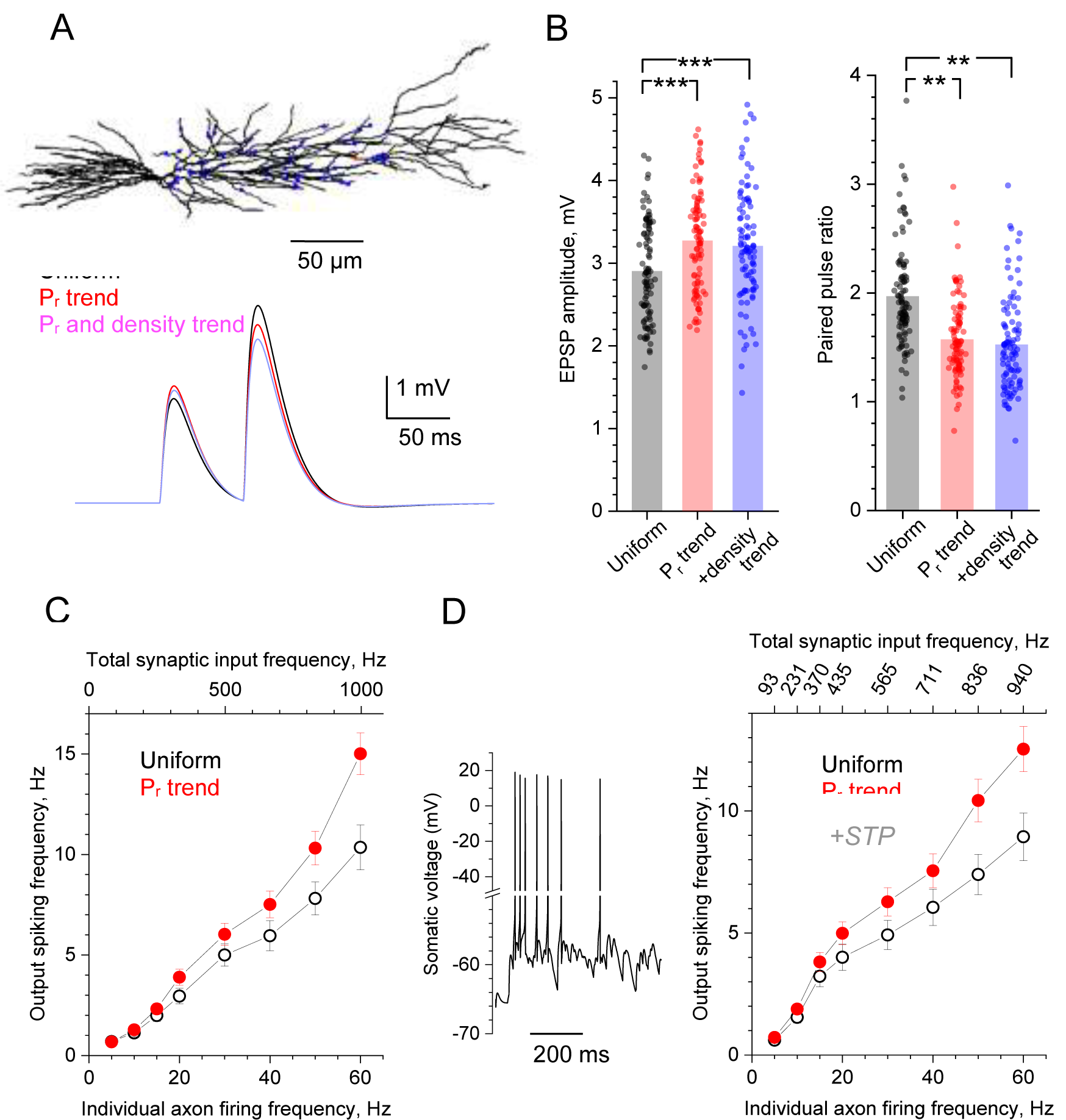
Computer simulations of CA1 pyramidal cell with stochastic excitatory synapses. (A) Diagram, NEURON model of a reconstructed CA1 pyramidal cell (Migliore et al., 1999) (ModelDB 2796; variable time step *dt*, t=34°C). 50 excitatory inputs (blue dots) generate bi-exponential conductance change (rise and decay time, 1 ms and 20 ms, respectively) stochastically, in accord with *P*_*r*_. Traces, simulated somatic response to paired-pulse stimuli (50 ms apart), with P_r_ distributed uniformly (black), in accord with the distance-dependent trend (as in Figure 1C; red), and both P_r_ and synaptic density trends (as in Figure 2E, blue). (B) Summary of test in (A); dots, individual runs (n = 100); bars, mean EPSP amplitude (left, mean ± SEM: 2.90 ± 0.060, 3.27 ± 0.061, 3.20 ± 0.068, for the three conditions, respectively, as indicated) and paired-pulse ratios (right, mean ± SEM: 1.96 ± 0.047, 1.57 ± 0.036, 1.52 ± 0.0437, notation as above) are shown; ***p < 0.005. (C) Input-output spiking rate relationship over the physiological range of input firing frequencies (per axon, bottom axis; total, top axis); hollow circles, uniform distribution of P_r_; solid symbols, P_r_ follows the distance-dependent trend (as in Figure 1C); mean ± SEM are shown (n = 100 simulation runs). (D) Trace: A characteristic cell spiking burst (model as in A) in response to a Poisson-process afferent spiking input (∼50 Hz per synapse) incorporating the experimental kinetics of short-term plasticity (STP) at CA3-CA1 synapses (Mukunda and Narayanan, 2017) (see Figure S4C-F for detail). Graph: Input-output spiking rate relationship across the physiological range of average input firing frequencies, with experimental STP incorporated; other notations as in (C); the top abscissa scale is nonlinear because STP affects average P_r_ in a biphasic, non-monotonous manner (see Figure S4C).

The results show that the P_r_ trend, on average, adds ∼13% to the single-pulse EPSP amplitude while decreasing PPR by ∼20%, compared to the evenly distributed P_r_, whereas adding the density trend produces little further change (Figure 4A-B).

Qualitatively similar results were obtained for the five-pulse burst tests based on the iGluSnFR data, suggesting that the P_r_ trend, when combined with the five-pulse slope trend (Figure S3C), boosts the voltage transfer value during the burst activity by ∼15% (Figure S4A-B).

However, the main role of P_r_ in synaptic signal integration and transfer is played out when a time series of afferent spikes, rather than one synchronous discharge, generate a postsynaptic spiking response (Thomson, 2000; Williams and Stuart, 2002; Williams and Atkinson, 2007; Grillo et al., 2018). Therefore, in the second test we simulated a Poisson process of stochastic synaptic discharges, across the physiological range of frequencies, and monitored the postsynaptic cell output. Experimental attempts to explore stochastic input to principal neurons have often employed synchronous (extracellular) activation of multiple afferent fibres, which is unlikely to happen in an intact brain where individual inputs can generate independent spike series. Thus, relatively intense synaptic input activity seen in pyramidal cells in vivo could correspond to relatively low spiking frequency at individual multiple afferents converging onto these cells (Bahner et al., 2011; Kowalski et al., 2016). These considerations suggest that short-term plasticity, if any, of intense synaptic input is actually driven mainly by the evolution of P_r_ values over low or moderate spiking frequencies at individual axons.

To understand whether and how the distance dependence of P_r_ influences spiking output of the postsynaptic pyramidal cell, we first compared the case of uniform P_r_ values (P_r_ = 0.36), with that of the P_r_ trend (the spine density trend was ignored as it had a negligible effect on EPSCs; Figure 4B). The outcome shows that having the P_r_ trend provides a clear advantage for signal transfer fidelity, which is particularly prominent at higher input frequencies (Figure 4C). However, in these simulations we assumed no use-dependent changes in the transmission efficacy as set by P_r_. In fact, a recent study of the CA3-CA1 circuitry has established the time-dependent degree of (presynaptic) short-term plasticity (STP) during afferent bursts, over a wide range of firing frequencies (Mukunda and Narayanan, 2017). In brief, during repetitive presynaptic activity, P_r_ undergoes prominent facilitation and depression within the first 2-4 discharges, followed by a near-constant P_r_ value that depends on the average input frequency (Mukunda and Narayanan, 2017). We have incorporated this STP algorithm (Figure S4C) and successfully validated it in our model by comparing simulated and recorded outcomes of the fixed-frequency afferent bursts (Figure S4D-F). Equipped with this STP mechanism, we generated Poisson-process afferent input in our model and found, once again, that the P_r_ trend improves signal transfer, with the effect monotonically increasing with higher frequencies of afferent firing (Figure 4D).

## DISCUSSION

The way information is handled and stored by brain circuits depends largely on the rules of synaptic signal integration by principal neurons. Passive electric properties of neuronal dendrites impose distance-dependent attenuation of synaptic receptor current. Thus, distal synapses should have a progressively weaker influence on somatic spike generation. If true, this would mean that neurons with large dendritic trees possess a highly inefficient synaptic connectome. However, studies in hippocampal CA1 pyramidal cells have found that synaptic receptor numbers increase with the synapse-soma distance (Andrasfalvy and Magee, 2001; Nicholson et al., 2006) and that ion channel properties at distal dendrites enable local regenerative events (Cash and Yuste, 1999; Magee, 1999). These features seem to underpin near-linear summation of synaptic inputs across the entire tree (Cash and Yuste, 1999; Magee, 2000; Magee and Cook, 2000), thus rescuing the efficiency of dendritic signal integration. However, equal contribution of individual axo-dendritic synapses to somatic signal assumes that their average discharge rates are similar. This in turn directly depends on P_r_ and its use-dependent plasticity at individual synapses (Williams and Stuart, 2002; Williams and Atkinson, 2007). It was therefore a surprising discovery that the P_r_ values at the basal dendrites of CA1 pyramidal cells showed a centrifugal decrease, at least in conditions of synchronous, multi-synaptic paired-pulse stimuli (Grillo et al., 2018). If common, this trend, again, would question the efficiency of synaptic connectome: it seems wasteful to make so many distal synaptic connections that have so little impact.

Here, we used OQAs to measure P_r_ at individual synapses in apical dendrites of CA1 pyramidal cells and found the opposite: P_r_ values were increasing with the synapse-soma distance along the dendrites. There are certain advantages of the OQA over paired-pulse experiments. Firstly, it establishes the actual P_r_ values and their distribution pattern along the dendrites. Secondly, the OQA deals with confirmed afferent activation of individual synapses and thus avoids the potential dependence between excitability of an axon and P_r_ value at its synapses, which would be masked in multiple-fibre stimulation. Thirdly, intrinsic differences in the short-term plasticity kinetics among different synaptic populations may contribute to the difference in PPRs. In addition, the outcome of the P_r_ assessment based on the use-dependent NMDA receptor blocker MK-801 (Grillo et al., 2018) may depend on the degree of extra-and cross-synaptic activation of NMDA receptors (Scimemi et al., 2004). The technically strenuous OQA has its own limitations, of which the main one is the low signal-to-noise ratio, at least at some connections. In addition, the high-affinity (slowly-unbinding) Ca_2+_ indicators required to detect dendritic spine signals tend to saturate rapidly, thus preventing reliable P_r_ readout for more than ∼2 successive stimuli. We therefore focused on synapses that showed a clear distinction between failures and successes during paired-pulse stimulation (Figure 1B), noting that this sampling still safely included the P_r_ range from 0.1-0.9 (Figure 1C). To further address a potential bias in selecting dendritic spines, we employed an alternative experimental design, in which P_r_ trends were assessed in the bulk of synapses in area CA1 by imaging evoked glutamate release with the genetically encoded optical sensor iGluSnFR. Similar to the earlier method (Grillo et al., 2018), we used PPR values to assess position-dependent variations in P_r_. While the earlier method referred to the position of the stimulation electrode with respect to the patched postsynaptic cell morphology, our method provided an exact spatial reference to the site of glutamate release in the *s. radiatum*.

We have found that both classical OQA and the iGluSnFR approach point to the centrifugal increase of P_r_ in apical dendrites of CA1 pyramidal cells. This lends support to the idea that synaptic organisation can powerfully compensate for the ‘electrotonic weakness’ of synapses in distal dendrites. The effects found here are similar to the location-dependent features of synaptic inputs revealed with paired recordings between L2/3 and L5 pyramidal neurons in the neocortex (Williams and Stuart, 2002; Williams and Atkinson, 2007). Clearly, the occurrence of higher-efficacy synapses at distal dendrites must boost temporal summation of local CA3-CA1 inputs (Stuart and Spruston, 2015). This is consistent with the observation that LTP induction there involves local sodium spikes (Kim et al., 2015). Furthermore, it has recently been found that individual CA3-CA1 axons make multiple synapses in distal (rather than proximal) dendrites of CA1 pyramidal cells (Bloss et al., 2018). This further argues for added efficiency of distal synaptic connections, and is consistent with the relatively high synaptic density found here. Because the spine density estimate could be affected by limited resolution of two-photon excitation imaging (which may not necessarily detect smallest or thinnest spines), it would be prudent to assume that here we refer mainly to larger, mushroom-type dendritic spines.

Notwithstanding methodological differences, comparing the present results with the earlier findings (Grillo et al., 2018) suggests that the dendritic integration traits at basal and apical inputs in CA1 pyramidal cells are starkly different. Excitatory inputs to basal dendrites in the *s. oriens* include axons of other pyramidal cells, septal fibre, and commissural axons from the contralateral hippocampus. Interestingly, the spine density here increases almost two-fold away from the soma (Bannister and Larkman, 1995; Ballesteros-Yanez et al., 2006). Whether and how these different inputs cluster within different parts of the basal dendrites, and how their spiking activities differ, might shed light on the adaptive role of the apparently ‘inefficient’ P_r_ pattern there. For instance, biophysical modelling of a simplified neuron suggested that the P_r_ and PPR trends in basal dendrites help to facilitate supra-linear summation of local generative events such as dendritic spikes (Grillo et al., 2018). Indeed, such events have long been considered as a key mechanism of distal dendritic signalling (Larkum and Nevian, 2008; Branco and Hausser, 2009).

In the present study, we explored a realistic model of a CA1 pyramidal cell (Migliore et al., 1999) hosting excitatory synapses that incorporated varied P_r_ values, and were scattered across the apical dendrites, in accord with our experimental observations. Simulations suggested that having the centrifugal P_r_ trend boosts synaptic signal transfer across the physiological frequency range, with the effect monotonically increasing with higher firing frequencies. Importantly, in an intact rat brain, CA1 pyramidal cells tend to fire in short, high-frequency bursts (Bahner et al., 2011; Kowalski et al., 2016) associated with sharp-wave ripple complexes (Bahner et al., 2011). The latter type of synchronised neuronal activity is thought to represent a powerful mechanism of memory consolidation in the hippocampus (Gruart et al., 2006; Buzsaki, 2015). In fact, short bursts of activity have long been considered as a key unit of information transfer by stochastic synapses (Lisman, 1997). Thus, the P_r_ trend found here in apical dendrites must play an essential role in supporting and enhancing the cellular underpinning of learning and memory in the brain.

## METHODS

### Hippocampal Slice Preparation

Animal procedures were subject to local ethical approval and adhered to the European Commission Directive (86/609/ EEC) and the United Kingdom Home Office (Scientific Procedures) Act of 1986. Male Sprague Dawley rats (P20-P28) were sacrificed using an overdose of isoflurane. Animals were kept in groups (5–8 per cage) under standard housing conditions with 12 h light–dark cycle and free access to food pellets and drinking water. Following decapitation brains were rapidly removed, hippocampi dissected out and transverse slices (350 µm thick) were prepared on a Leica VT1200S vibratome. The slicing was performed in ice-cold slicing solution that contained (in mM): 60 NaCl, 105 sucrose, 26 NaHCO_3_, 2.5 KCl, 1.25 NaH_2_PO_4_, 7 MgCl_2_, 0.5 CaCl_2_, 11 glucose, 1.3 ascorbic acid and 3 sodium pyruvate. Alternatively, slices were cut in an ice-cold slicing solution that contained (in mM): 64 NaCl, 2.5 KCl, 1.25 NaH_2_PO_4_, 0.5 CaCl_2_, 7 MgCl_2_, 25 NaHCO_3_, 10 D-glucose, and 120 sucrose. All solutions were bubbled with 95% O_2_ plus 5% CO_2_, pH adjusted to 7.4. Once cut, slices were incubated in the oxygenated slicing solution at 34 °C for 15 min and then allowed to equilibrate to room temperature for 15 min following which they were transferred to either a continuously oxygenated humid interface chamber, or a submersion chamber. Both chamber types contained an oxygenated artificial cerebrospinal fluid (aCSF) solution containing (in mM): 120 NaCl, 10 glucose, 2.5 KCl, 1.3 MgSO_4_, 1 NaH_2_PO_4_, 25 NaHCO_3_, 1.3 MgCl_2_, 2 CaCl_2_, with an osmolality of ∼300 mOsm. Slices were then allowed to recover at room temperature for 1-5 hours prior to being transferred to the recording chamber of the microscope constantly perfused with 32–34 °C.

### Electrophysiology *ex vivo*

Whole-cell patch-clamp recordings were performed from CA1 pyramidal cells visualised using infrared differential contrast imaging. Thin-walled borosilicate glass capillaries were used to fabricate recording electrodes with a resistance of 2.5–3.5 MΩ. Intracellular pipette solution contained (in mM) KCH_3_O_3_S 135, HEPES 10, Na_2_-Phosphocreatine or di-Tris-Phosphocreatine 10, MgCl_2_ 4, Na_2_-ATP 4, Na-GTP 0.4, 5 QX-315-Bromide (pH adjusted to 7.2 using KOH, osmolarity 290-295). Cell-impermeable Ca^2+^ dyes detailed below and the Ca^2+^ insensitive morphological tracer Alexa Fluor 594 hydrazide (50 μM) were routinely added to the intracellular solution. Throughout recordings cells were held in voltage clamp at -65mV, and the aCSF routinely supplemented with 100µM Picrotoxin (Tocris Bioscience) and 30µM D-Serine. Electrophysiological recordings were carried out by using a set of remotely controlled micromanipulators and XY-translation stage (Luigs and Neumann) with a Multiclamp 700B amplifier (Molecular Devices). Signals were digitized at 10 kHz and stored for off-line analysis using WinWCP V4.1.5 -4.7 (John Dempster, University of Strathcyde) or PClamp 10.4-10.5 (Molecular Devices)

### Two-photon excitation imaging: optical quantal analysis

We used a Radiance 2100 imaging system (Zeiss–Bio-Rad; 60x Olympus objective, NA0.9) or a Femtonics Femto3D-RC imaging system (25x Olympus objective, NA1.05) both optically linked to two femtosecond pulse lasers MaiTai SpectraPhysics-Newport) and integrated with patch-clamp electrophysiology.

Intracellular Ca^2+^ responses were routinely imaged in the Ca^2+^ sensitive (green) emission channel filtered to the bandwidth of 500-550nm, with either of the Ca^2+^ sensitive dyes Oregon Green BAPTA-1 (250 μM, Invitrogen), Fluo-4 (400 μM, Invitrogen) or Fluo-8 (300 µM, Stratech Scientific). For optical quantal analysis synaptically evoked responses in dendritic spines were triggered by minimal electrical stimulation using a monopolar glass stimulating electrode filled with aCSF and placed 10-20 µm from the dendritic branch targeted. To identify active synapses relatively fast (10 Hz) frame scans of the local dendrites were viewed whilst three 100 μs square pulses of 2-10 V were delivered with a 25 ms inter-stimulus interval using a constant voltage isolated stimulator (model DS2A-mkII, Digitimer Ltd, Welwyn Garden City, UK). This protocol was repeated until a Ca^2+^ response confined to a spine head was observed, then 400-700 ms line scans of the active spine were then recorded at a line scan rate of 500 Hz. In order to reduce the inherent bias toward high release probability synapses a dual stimulus protocol (50 ms inter-stimulus interval) was employed to ensure response stability at synapses with low initial release probability. Line scans were repeated once every 30 s with a minimum of 15 trials to assess release probability and manually corrected for focus fluctuations in the Alexa (red) channel. Line scan profiles were extracted using ImageJ 1.x (Schneider et al., 2012) or Femtonics MES 6 and routinely documented as *ΔG/R* where *ΔG* = *G - G*_*0*_ stands for the fluorescence signal in the green channel *G* with the baseline fluorescence *G*_*0*_ (averaged over the baseline time window) subtracted, and *R* stands for the Alexa fluorescence in the red channel (corrected for photobleaching, if any). Where recordings of synaptic responses were made at two sites along a single dendritic branch (distal and proximal to the branch point), equal numbers of recordings first proximal – distal and distal - proximal were obtained to remove bias in the temporal order by which the recordings were carried out. Experiments were excluded if the synapse became non-responsive, if spine morphology visibly changed during recording or if significant drift of the stimulation electrode occurred. Release probability (P_r_) was defined as the probability of success on the first stimulus (shown as P1 in figures, sum of successful trials/total number of trials). Where a second response was clearly visible over the first, P2 was also measured and defined as the sum of responses to the second stimulus + responses to both stimuli/total number of trials. PPR was defined as P2/P1 = P2/P_r_.

### Two-photon excitation imaging: Morphological measurements

For morphological tracing purposes, wide field high resolution images of the apical dendritic tree (>75 µm^2^) were acquired at the maximal optical resolution as a series of individual 3-D stacks (50-100 μm deep each) in the Alexa emission channel (550-650 bandwidth; λ_x_^2p^ = 800 nm), collected in image frame mode (Biorad: 512 × 512 pixels, 8-bit; Femto3D >512×512 pixels, 16-bit) at 1.5-2.5 µm steps. These images were used to determine the location of the individual synapses in relation to the CA1 pyramidal cell soma, first the averaged Alexa 594 image stack was used to form a template defining the two-dimensional x-y path (along the dendrite) between soma and spine of interest. The start point of the distance measurement was defined as the centre of an oval shaped to fit the edges of the imaged soma, from this point the segmented line function in ImageJ was used to trace the path along the dendritic tree between the soma and the spine of interest ending at the base of the spines shaft. This line was then saved as a region of interest and then overlaid onto the original 3D image stack. The ‘reslice’ function in ImageJ was then used to form a two-dimensional y-z image in which the y-axis represents the length of the template line and the z-axis representing the deviation in the depth of the slice thus enabling simple measurement of distance from soma to synapse with z-plane deviation taken into account. Using this method distances from the origin of and to the end of the parent dendrite were also obtained. Higher magnification images of the same resolution were also acquired and the density of spines local to the synapse estimated by counting the total number of visible protrusions along a 10 µm portion of dendrite centred on the synapse of interest.

### Viral transduction for labelling Schaffer collateral axonal boutons

For ex vivo imaging of axonal boutons, viral transduction in vivo via neonatal intracerebroventricular (ICV) injections in both male and female C57BL/6J mice (Charles River Laboratories) were used, as we detailed earlier (Kopach et al., 2020). Briefly, an AAV virus expressing the neuronal optical glutamate sensor, AAV9.hSynap.iGluSnFR.WPRE.SV40 (supplied by Penn Vector Core, PA) was injected into the cerebral ventricles of neonates (P0-P1) during aseptic surgery. The viral particles were injected at 2 µl per hemisphere (2.5-5 × 10^9^ genomic copies in total), at a rate not exceeding of 0.2 µl/s, 2 mm deep, guided to a location approximately 250 µm lateral to the sagittal suture and 500–750 µm rostral to the neonatal coronary suture. After animals received AAV injections, they were returned to the mother in their home cage; they were systematically kept as a group of litters and monitored for days thereafter, to ensure that no detrimental side effects appear. Satisfactory transduction of iGluSnFR *in vivo* occurred within 3-4 weeks.

### Two-photon excitation (2PE) imaging of glutamate release with iGluSnFR

iGluSnFR fluorescence was recorded in the green emission channel under 2PE at λ_x_^2P^=910 nm, using a Femtonics Femto3D-RC imaging system or an Olympus FV10MP imaging system, both optically linked to a Ti:Sapphire MaiTai femtosecond-pulse laser (SpectraPhysics-Newport), equipped with galvo scanners, and integrated with patch-clamp electrophysiology. In the *s*.*radiatum*, we focused on individual axonal boutons that could be visualised by iGluSnFR expression and showed a consistent optical response to electric stimuli applied to Schaffer collaterals. To minimise photodamage, only a single focal section through the region of interest (ROI) containing selected axonal fragments was imaged, at laser power not exceeding 3-6 mW under the objective. The focal plane was regularly adjusted, to account for specimen drift. To record optical signals with high spatiotemporal resolution while minimising photodamage, we employed the scanning mode of spiral (‘tornado’) linescans centred at the bouton of interest, as detailed previously (Jensen et al., 2017; Jensen et al., 2019). In these experiments, we recorded responses to paired-pulse stimuli at 20 Hz, normally collecting 20-30 trials ∼30 s apart, and documented the shortest distance between the recorded bouton and the *s. pyramidale* border. The iGluSnFR fluorescence response to afferent stimulation was expressed as (*F*(*t*)*-F*_0_) */ F*_0_= *ΔF / F*_0_, where *F*(*t*) stands for fluorescence intensity over time, and *F*_0_ is the baseline intensity averaged over ∼150 ms prior to the first stimulus.

In a complementing experiment, we applied five-pulse stimuli (at 20 Hz) to Schaffer collaterals and used time-lapse imaging in frame-scan mode, as detailed previously (Kopach et al., 2020), thus providing short pixel dwell time (∼0.5 µs) to enable signal registration over larger imaged regions, albeit at the expense of temporal resolution. This approach was chosen because regions of interest had to be selected off-line post hoc, based on their overall morphological stability: burst stimulation could lead to microscopic focal drifts or tissue changes in some areas, which had to be avoided. Therefore, the experimental time course of iGluSnFR featuring a limited number of time points was fitted with the underlying theoretical kinetics of five overlapping *ΔF / F*_0_ signals so that: Δ*F* / *F*_0_ = ∑ *A* _*i*_ exp(−(*t* − Δ*t* = (*i* − 1)) · *τ* ^*−*1^) (*i* = 1,…, 5) where *A*_*i*_ is the *i*th signal amplitude (fitted directly to the recoded amplitude), *Δt* = 50 ms (inter-spike interval), and the decay constant *τ* obtained from fitting the signal decay (tail) after the fifth pulse.

### NEURON modelling

We employed a multi-compartmental NEURON (Hines and Carnevale, 2001) model of a reconstructed CA1 pyramidal neuron, with a full set of experiment-adjusted membrane currents, uploaded from the NEURON 7.6×64 database (https://senselab.med.yale.edu/modeldb; models 2796 and 7509). Simulations were performed using a variable time step *dt*; the cell axial specific resistance *R*_*a*_ and capacitance C_m_ were set at *R*_*a*_ = 90 Om·cm, and *C*_*m*_ = 1 mF/cm^2^, throughout simulations. Fifty synapses were distributed over the dendritic apical tree, between 10 and 350 µm from the soma, either uniformly randomly or in accord with the experimental statistical trend, as specified. Computation tests for any set of parameters routinely consisted of 100 runs (trials). To avoid any bias arising from a particular (fixed) distribution of synapses along the dendrites, this distribution was generated anew at each trial, albeit with the same probability density function. However, in tests where non-uniform and uniform distributions of Pr were compared, the random generator seed was kept unchanged.

At individual synapses, excitatory synaptic conductance *g*_*s*_*(t)* was modelled using the dual-exponential formalism realised in the NEURON function Exp2Syn. The dynamics models synaptic activation as a change in synaptic conductance with a time course given by: *g*_*s*_ = *G*_*m*_ =exp(−*t* / *τ*_1_) − exp(−*t* / *τ*_2_) where *τ*_*1*_ and *τ*_*2*_ are the rise and decay time constants, respectively, and *G*_*m*_ is the factor defined by the peak synaptic conductance. The reverse potential of excitatory synapses was zero, in line with that for AMPMA and NMDA receptor types. In line with the data reporting that synaptic strength increases centrifugally in apical dendrites of CA1 pyramidal cells (Andrasfalvy and Magee, 2001; Nicholson et al., 2006), *G*_*m*_ values were set proportional to the function *f(x)* = *A·*(0.51 + 0.002*x*) where *x* is the synapse-soma distance, and *A* is scaling factor (NEURON NetCon object), which in our case sets single-synapse conductance *G*_*m*_ (in µS). In simulations exploring subthreshold postsynaptic excitation, *A* was set at 0.002 µS, and in tests exploring the spiking input - output relationship *A* was set at 0.06 µS.

To replicate our experimental observations of release probability, the average P_r_ (P1) was set at 0.36, and P2 at 0.77. These values remained constant throughout the synaptic population in the case of the uniform P_r_ distribution. In the case of the experimental (centrifugal) P_r_ trend, the P_r_ (P1) and P2 values were set as a function of the synapse-soma distance *x*: P1 = (0.18 + 0.0012*x*), and P2 = (0.18 + 0.0012*x*)(3.31 - 0.0074*x*), reflecting experimental regressions. Calculations started 300 ms following full stabilisation of the NEURON model.

#### ACKNOWLEDGEMENTS

This study was supported by the Wellcome Trust Principal Fellowship (212251_Z_18_Z), ERC Advanced Grant (323113) and European Commission NEUROTWIN grant (857562).

## AUTHORS CONTRIBUTIONS

D.A.R. and T.P.J. conceived the study and its research strategies; T.P.J. designed and carried out electrophysiological and Ca^2+^ imaging experiments and analyses; O.K. carried out iGluSnFR transfections in neonates, iGluSnFR imaging experiments and analyses; J.P.R. established and optimised original iGluSnFR transfection protocols; L.P.S. designed and carried out biophysical simulations; D.A.R. carried out data analyses and wrote the manuscript, which was subsequently complemented by all authors.

## REFERENCES

Andrasfalvy BK, Magee JC (2001) Distance-dependent increase in AMPA receptor number in the dendrites of adult hippocampal CA1 pyramidal neurons. J Neurosci 21:9151–9159.

Araya R, Eisenthal KB, Yuste R (2006) Dendritic spines linearize the summation of excitatory potentials. Proc Natl Acad Sci U S A 103:18799–18804.

Bahner F, Weiss EK, Birke G, Maier N, Schmitz D, Rudolph U, Frotscher M, Traub RD, Both M, Draguhn A (2011) Cellular correlate of assembly formation in oscillating hippocampal networks in vitro. Proc Natl Acad Sci U S A 108:E607–616.

Ballesteros-Yanez I, Benavides-Piccione R, Elston GN, Yuste R, DeFelipe J (2006) Density and morphology of dendritic spines in mouse neocortex. Neurosci 138:403–409.

Bannister NJ, Larkman AU (1995) Dendritic morphology of CA1 pyramidal neurones from the rat hippocampus: II. Spine distributions. J Comp Neurol 360:161–171.

Bloss EB, Cembrowski MS, Karsh B, Colonell J, Fetter RD, Spruston N (2018) Single excitatory axons form clustered synapses onto CA1 pyramidal cell dendrites. Nat Neurosci 21:353–363.

Boddum K, Jensen TP, Magloire V, Kristiansen U, Rusakov DA, Pavlov I, Walker MC (2016) Astrocytic GABA transporter activity modulates excitatory neurotransmission. Nat Commun 7:13572.

Branco T, Hausser M (2009) The selfish spike: local and global resets of dendritic excitability. Neuron 61:815–817.

Branco T, Hausser M (2011) Synaptic integration gradients in single cortical pyramidal cell dendrites. Neuron 69:885–892.

Branco T, Staras K, Darcy KJ, Goda Y (2008) Local dendritic activity sets release probability at hippocampal synapses. Neuron 59:475–485.

Buzsaki G (2015) Hippocampal sharp wave-ripple: A cognitive biomarker for episodic memory and planning. Hippocampus 25:1073–1188.

Cash S, Yuste R (1999) Linear summation of excitatory inputs by CA1 pyramidal neurons. Neuron 22:383–394.

Emptage NJ, Reid CA, Fine A, Bliss TV (2003) Optical quantal analysis reveals a presynaptic component of LTP at hippocampal Schaffer-associational synapses. Neuron 38:797–804.

Grillo FW, Neves G, Walker A, Vizcay-Barrena G, Fleck RA, Branco T, Burrone J (2018) A Distance-Dependent Distribution of Presynaptic Boutons Tunes Frequency-Dependent Dendritic Integration. Neuron 99:275–282 e273.

Gruart A, Munoz MD, Delgado-Garcia JM (2006) Involvement of the CA3-CA1 synapse in the acquisition of associative learning in behaving mice. J Neurosci 26:1077–1087.

Harris KM, Stevens JK (1989) Dendritic spines of CA 1 pyramidal cells in the rat hippocampus: serial electron microscopy with reference to their biophysical characteristics. J Neurosci 9:2982–2997.

Hines ML, Carnevale NT (2001) NEURON: a tool for neuroscientists. Neuroscientist 7:123–135.

Jensen TP, Zheng K, Tyurikova O, Reynolds JP, Rusakov DA (2017) Monitoring single-synapse glutamate release and presynaptic calcium concentration in organised brain tissue. Cell Calcium 64:102–108.

Jensen TP, Zheng KY, Cole N, Marvin JS, Looger LL, Rusakov DA (2019) Multiplex imaging relates quantal glutamate release to presynaptic Ca2+ homeostasis at multiple synapses in situ. Nature Communications 10:1414.

Katz Y, Menon V, Nicholson DA, Geinisman Y, Kath WL, Spruston N (2009) Synapse distribution suggests a two-stage model of dendritic integration in CA1 pyramidal neurons. Neuron 63:171–177.

Kim Y, Hsu CL, Cembrowski MS, Mensh BD, Spruston N (2015) Dendritic sodium spikes are required for long-term potentiation at distal synapses on hippocampal pyramidal neurons. Elife 4.

Kopach O, Zheng KY, Rusakov DA (2020) Optical monitoring of glutamate release at multiple synapses in situ detects changes following LTP induction. Molecular Brain 13.

Kowalski J, Gan J, Jonas P, Pernia-Andrade AJ (2016) Intrinsic membrane properties determine hippocampal differential firing pattern in vivo in anesthetized rats. Hippocampus 26:668–682.

Larkum ME, Nevian T (2008) Synaptic clustering by dendritic signalling mechanisms. Curr Opin Neurobiol 18:321–331.

Lisman JE (1997) Bursts as a unit of neural information: making unreliable synapses reliable. Trends Neurosci 20:38–43.

Magee JC (1999) Dendritic lh normalizes temporal summation in hippocampal CA1 neurons. Nat Neurosci 2:508–514.

Magee JC (2000) Dendritic integration of excitatory synaptic input. Nat Rev Neurosci 1:181–190.

Magee JC, Cook EP (2000) Somatic EPSP amplitude is independent of synapse location in hippocampal pyramidal neurons. Nat Neurosci 3:895–903.

Marvin JS et al. (2018) Stability, affinity, and chromatic variants of the glutamate sensor iGluSnFR. Nat Methods 15:936–939.

Migliore M, Hoffman DA, Magee JC, Johnston D (1999) Role of an A-type K+ conductance in the back-propagation of action potentials in the dendrites of hippocampal pyramidal neurons. J Comput Neurosci 7:5–15.

Mukunda CL, Narayanan R (2017) Degeneracy in the regulation of short-term plasticity and synaptic filtering by presynaptic mechanisms. J Physiol 595:2611–2637.

Nicholson DA, Trana R, Katz Y, Kath WL, Spruston N, Geinisman Y (2006) Distance-dependent differences in synapse number and AMPA receptor expression in hippocampal CA1 pyramidal neurons. Neuron 50:431–442.

Oertner TG, Sabatini BL, Nimchinsky EA, Svoboda K (2002) Facilitation at single synapses probed with optical quantal analysis. Nature Neurosci 5:657–664.

Rusakov DA, Kullmann DM (1998) Extrasynaptic glutamate diffusion in the hippocampus: ultrastructural constraints, uptake, and receptor activation. J Neurosci 18:3158–3170.

Schneider CA, Rasband WS, Eliceiri KW (2012) NIH Image to ImageJ: 25 years of image analysis. Nat Methods 9:671–675.

Scimemi A, Fine A, Kullmann DM, Rusakov DA (2004) NR2B-containing receptors mediate cross talk among hippocampal synapses. J Neurosci 24:4767–4777.

Stuart GJ, Spruston N (2015) Dendritic integration: 60 years of progress. Nat Neurosci 18:1713–1721.

Sylantyev S, Jensen TP, Ross RA, Rusakov DA (2013) Cannabinoid-and lysophosphatidylinositol-sensitive receptor GPR55 boosts neurotransmitter release at central synapses. Proc Natl Acad Sci USA 110:5193–5198.

Thomson AM (2000) Molecular frequency filters at central synapses. Progr Neurobiol 62:159–196.

Williams SR, Stuart GJ (2002) Dependence of EPSP efficacy on synapse location in neocortical pyramidal neurons. Science 295:1907–1910.

Williams SR, Atkinson SE (2007) Pathway-specific use-dependent dynamics of excitatory synaptic transmission in rat intracortical circuits. J Physiol 585:759–777.

Zador A (1998) Impact of synaptic unreliability on the information transmitted by spiking neurons. J Neurophysiol 79:1219–1229.

Zheng K, Bard L, Reynolds JP, King C, Jensen TP, Gourine AV, Rusakov DA (2015) Time-resolved imaging reveals heterogeneous landscapes of nanomolar Ca2+ in neurons and astroglia. Neuron 88:277–288.

