## Supplemental Figures for "Release probability increases towards distal dendrites boosting high-frequency signal transfer"

### Supplementary Figures

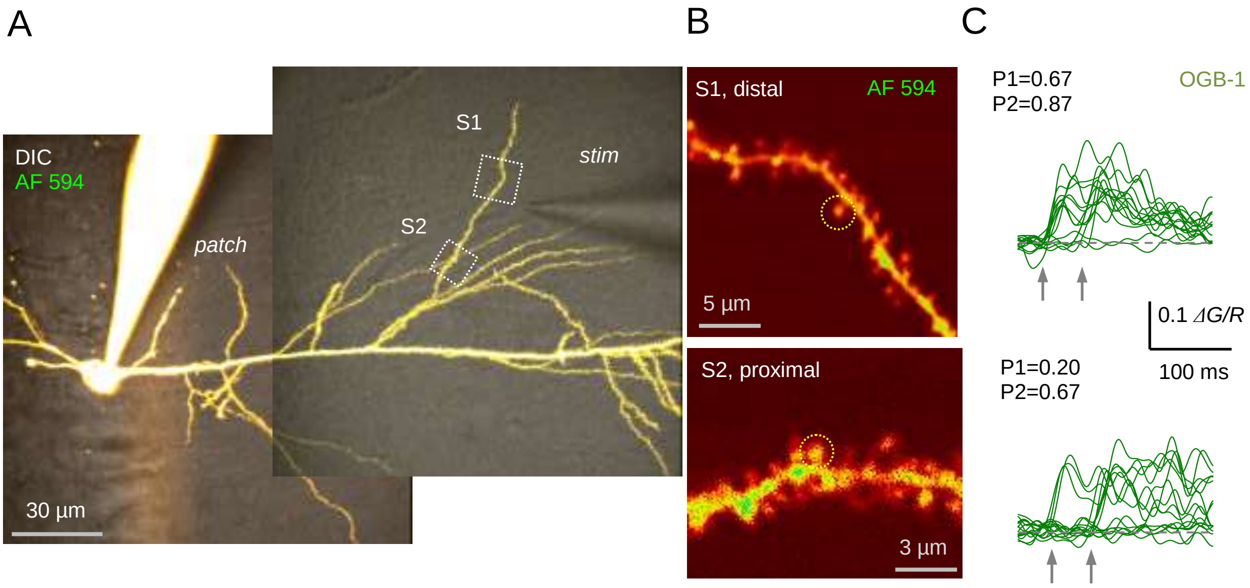


**Figure S1. Optical quantal analysis at individual CA3-CA1 synapses in acute hippocampal slices: second-order dendrite.**

(A) CA1 pyramidal cell held in whole-cell mode dialysed with 50 µM AF 594 and 400 µM OGB-1 (75 µm z-stack average, $\lambda_x^{2p}$ = 800 nm; DIC + AF 594 channel combined). Patch pipette (patch) and stimulating electrode (stim) can be seen; dotted rectangles (S1 and S2), ROIs to record from individual dendritic spines.

(B) ROIs as shown in (A) by dotted, at higher magnification (AF 594 channel only); dotted circles, two dendritic spines of interest.

(C) Examples of $Ca^{2+}$ linescan signal traces ($\Delta G/R$, green-channel OGB-1 increment signal $\Delta G$ related to red-channel AF 594 signal $R$) recorded in two dendritic spines as shown in (B), in response to two afferent stimuli (arrows) applied by a stimulating electrode (as in A). Release failures and successes can be clearly separated; P1 and P2, average release probability in response to the first and second stimulus, respectively.

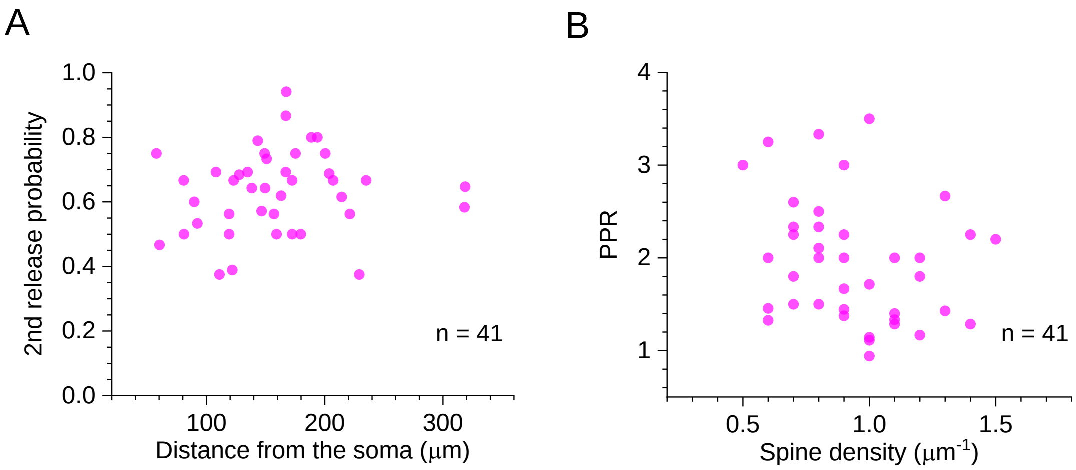


**Figure S2. Selected features of excitatory synapses with respect to their dendritic location.**

(A) Average probability of the second release (paired-pulse stimuli 50 ms apart) plotted against distance to the soma; sample mean ± SEM: 0.633 ± 0.20 (n = 41).

(B) Paired-pulse ratio (as P2/P1 in Supplementary Figure 1C) plotted against spine density along dendrites; sample mean ± SEM: 2.14 ± 0.17.

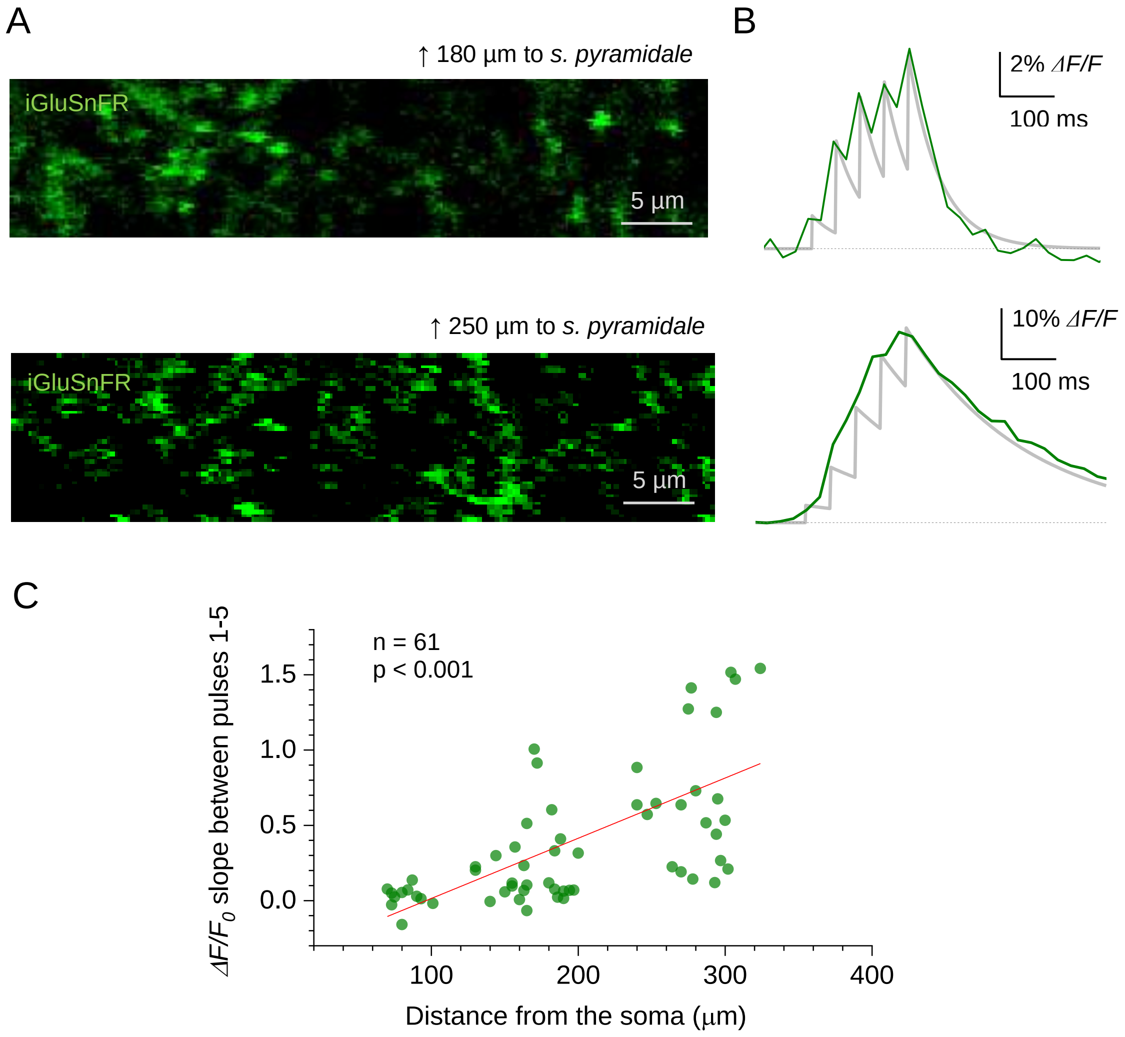


**Figure S3.** Optical (multi-synaptic) glutamate signal evoked by short bursts of Schaffer collateral stimulation at varied distances from *s. pyramidale*.

(A) Examples of recorded ROIs (iGluSnFR green fluorescence channel) in *s. radiatum*, with groups of tentative axonal boutons expressing iGluSnFR, at two different distances to the *s. pyramidale* border (as illustrated in Figure 3A; distance counted to the ROI horizontal midline).

(B) ROI-average iGluSnFR responses (green line, temporal resolution ~25 ms) to five stimuli 50 ms apart. The underlying sensor signal kinetics (light grey line ) was reconstructed using the fitting algorithm $\Delta F / F_0 = \sum A_i \exp\left(-\left(t - \Delta t \cdot (i-1)\right) \cdot \tau^{-1}\right)$ ($i$ = 1,..., 5) where $A_i$ is the $i$th signal amplitude (fitted directly to the recoded amplitude), $\Delta t$ = 50 ms, and the decay constant $\tau$ obtained from fitting the signal tail after the fifth pulse.

(C) Average slope (linear regression value) of the $\Delta F/F_0$ responses between the onsets for the 1st and 5th stimuli, at different distances from the *s. pyramidale*. (Pearson's r = 0.641).

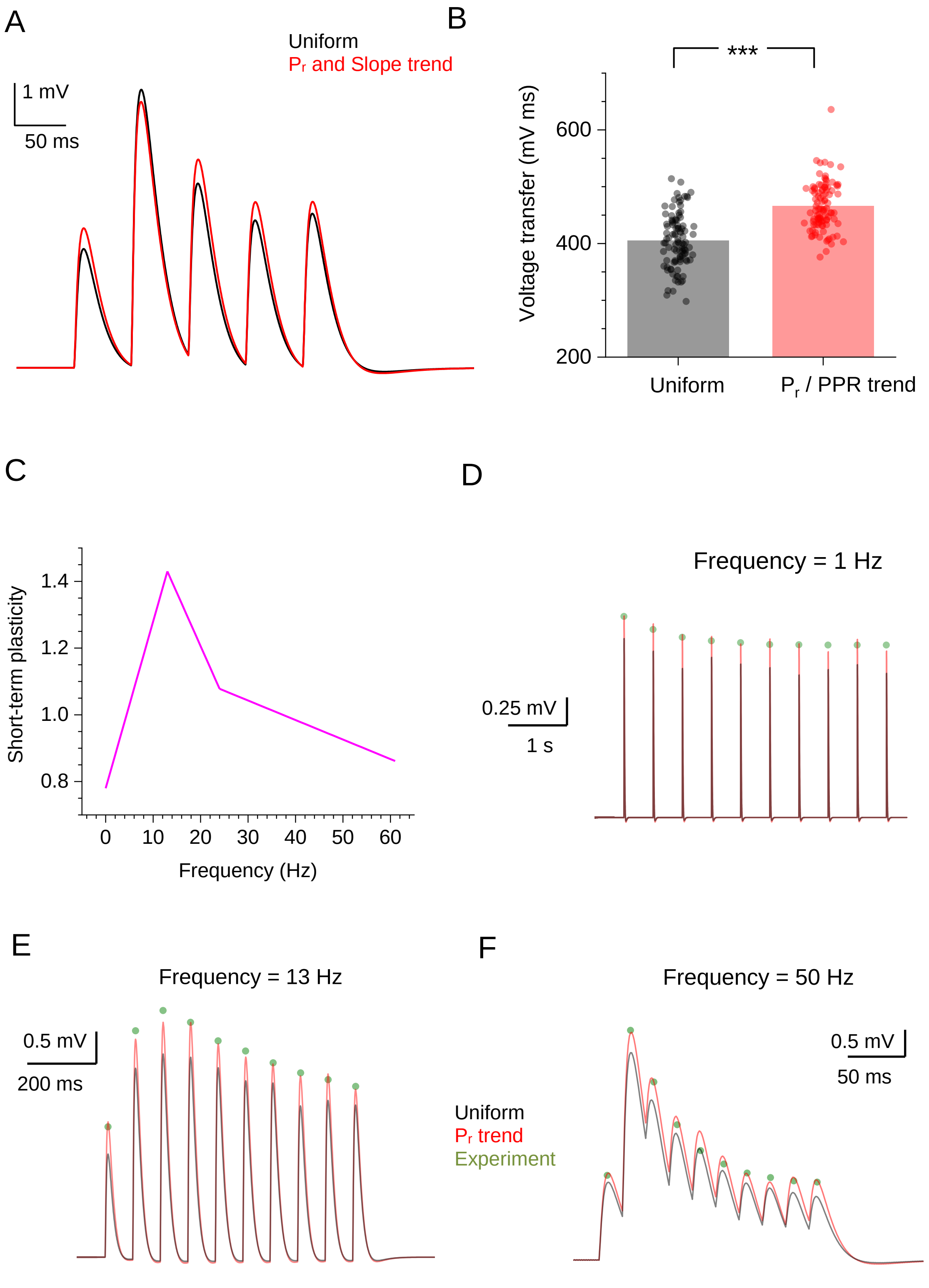


**Figure S4. Simulating the experiment-based kinetics of short-term plasticity (STP) in the CA3-CA1 circuit.**

(A) Traces show simulated average somatic response to five-burst 20 Hz stimuli (NEURON model as in Figure 4A), with synaptic Pr values distributed uniformly (black line), and in accord with the distance-dependent trend of $P_r$ (as in Fig. 1C-D) and the five-pulse $\Delta F/F_0$ slope (as in Fig. S3C).

(B) Summary of stochastic simulations shown in (A); ordinate, voltage transfer (area under the voltage curve over the five EPSPs); dots, individual runs (n = 100); bars, average values; ***p < 0.001.

(C) Linearized representation of the STP ratio profile for the 1–50 Hz range of presynaptic spiking frequencies replicating the experimental band-pass structure of the CA3–CA1 Schaffer collateral synapses (1).

(D-F) Simulated EPSP response in a CA1 pyramidal cell (NEURON model as in Figure 4A) to a burst of afferent stimuli, with synaptic $P_r$ values distributed uniformly (black), and in accord with the distance-dependent trend of $P_r$ (as in Figure 1C; red); green dots, experimental data (from (1)), normalised to the first EPSP amplitude, as indicated; postsynaptic response incorporates STP kinetics, as shown in (C).

### Supplementary references

1. Mukunda CL & Narayanan R (2017) Degeneracy in the regulation of short-term plasticity and synaptic filtering by presynaptic mechanisms. *J Physiol* 595(8):2611-2637.
